# Comparative connectomics reveals stage-specific gap junction rewiring that reshapes avoidance behavior

**DOI:** 10.64898/2026.05.28.728331

**Authors:** Daniel T. Choe, Ken C.Q. Nguyen, Myunghwan Choi, David H. Hall, J. Alexander Bae, Junho Lee

## Abstract

Environmental stress can remodel neural circuits, yet how such rewiring generates behavioral adaptation remains poorly understood. Here, we investigated how dauer-specific circuit remodeling alters nociceptive behavior. We discovered that dauers exhibit significantly shorter avoidance durations than adults. Through comparative connectomics, we observed a substantial expansion of gap junctions within the dauer nociceptive circuit and found that dauer neurons exhibit more transient activity dynamics as a result of this rewiring. Modeling neuronal activity demonstrated that removing dauer-specific connectivity constraints failed to reproduce transient characteristics, validating the functional role of dauer-specific gap junctions. Furthermore, introducing synthetic dauer-specific gap junctions into adults was sufficient to recapitulate dauer-like activity patterns and shorten avoidance duration. Despite extensive rewiring, avoidance initiation remained preserved across developmental stages. Our findings show that the dauer neural circuit is selectively rewired through gap junction remodeling to balance flexibility and stability for survival under stress.

## INTRODUCTION

Understanding how differences in neural circuitry give rise to differences in behavior is a central question in neuroscience. *Caenorhabditis elegans* (*C*. *elegans*) provides a uniquely powerful system for addressing this problem, as it remains the only organism for which connectomes have been reconstructed across all major developmental stages and both sexes (*1–4*). Among these stages, the dauer represents an alternative developmental form entered under adverse environmental conditions such as food scarcity or elevated temperature (*5*, *6*). The dauer stage is associated with profound behavioral adaptations that distinguish it from reproductive stages. For example, dauers are attracted to CO_2_ (*7*), whereas adults typically avoid it (*8*, *9*); they exhibit faster locomotion (*5*, *10*); and they display nictation, in which worms raise and wave their bodies in three dimensions (*11*). These qualitative behavioral shifts raise a fundamental question: how does dauer-specific circuit rewiring support stage-specific survival strategies?

The recent reconstruction of the dauer chemical synapse connectome has enabled systematic investigation of how neural circuits are developmentally remodeled under stress conditions and how such remodeling may shape behavior (*4*, *12*). Network-level analyses revealed that while interneuron-centered connectivity remains largely stable across development, the outputs of sensory neurons are extensively rewired in dauers (*4*). These findings suggest that stage-specific sensory processing may be a major driver of dauer-specific behavioral adaptation and raise the possibility that specific sensory circuits are selectively modified to tune behavioral outputs for survival during developmental arrest.

In addition to chemical synapses, neural circuits are connected by electrical synapses, or gap junctions, which permit direct current flow between neurons and enable rapid signal transmission (*13–15*). Electrical synapses are increasingly recognized as dynamic regulators of neural circuit function that contribute to synchronization, state transitions, sensory processing, locomotion, and behavior across diverse nervous systems (*13*, *16*). Indeed, studies in both vertebrates and invertebrates have demonstrated that electrical synapses can shape motor outputs, locomotor state transitions, and escape behaviors by regulating the timing and coordination of neuronal activity (*16–18*). In *C*. *elegans*, gap junctions formed by innexin proteins are widespread throughout the nervous system and play important roles in locomotion, sensory processing, and circuit function (*19*). These observations raise the possibility that developmental remodeling of electrical synapses contributes to the emergence of dauer-specific behaviors.

To determine how dauer-specific circuit remodeling influences behavior, we focused on the nociceptive circuit, in which ASH sensory neurons relay aversive signals to downstream command interneurons (AVA, AVB, AVD, and PVC), which in turn engage A- and B-type motor neurons to generate escape behavior (*20–24*). In *C*. *elegans*, nociceptive stimuli evoke a stereotyped avoidance response in which animals initiate backward locomotion, reorient, and subsequently resume forward movement. This behavior can be conceptually divided into two components: avoidance initiation, which determines whether an animal begins a reversal in response to a stimulus, and avoidance duration, which determines how long the reversal is maintained before forward locomotion resumes. These two components define the stochastic and temporal structure of nociceptive escape behavior, and separating them allows circuit organization to be related to distinct features of behavioral dynamics. Given the extensive sensory rewiring observed in the dauer connectome, we asked how these components of nociceptive behavior are altered in dauers and how such changes relate to circuit organization and activity. Here, we show that dauer-specific gap junctions shorten avoidance duration while leaving avoidance initiation intact, revealing a form of developmental rewiring that reshapes the timing of escape behavior without compromising its reliability.

## RESULTS

### Dauers exhibit shortened avoidance duration compared with adults

Freely moving *C*. *elegans* moves through sequences of forward movement, reversals, and turns, allowing them to avoid aversive stimuli and navigate toward favorable cues such as food (Fig. 1A). Upon detection of a nociceptive signal by ASH sensory neurons, *C*. *elegans* responds through a stereotyped escape sequence in which animals abruptly halt forward locomotion, execute a reversal, and reorient away from the aversive source (Fig. 1A, inset). We divide this escape behavior into two components: 1) avoidance initiation, whether an animal begins a reversal, and 2) avoidance duration, how long the reversal is maintained before forward locomotion resumes. Since nociceptive avoidance is stochastic, initiation is quantified as the probability of reversal across repeated trials (*23*). We first focused on avoidance duration, as differences between dauer and adult animals were immediately apparent. Notably, adult animals appeared to sustain longer reversal behaviors than dauers (Fig. 1B,C), and we therefore quantitatively compared the avoidance duration between the two stages.

**Fig. 1.**
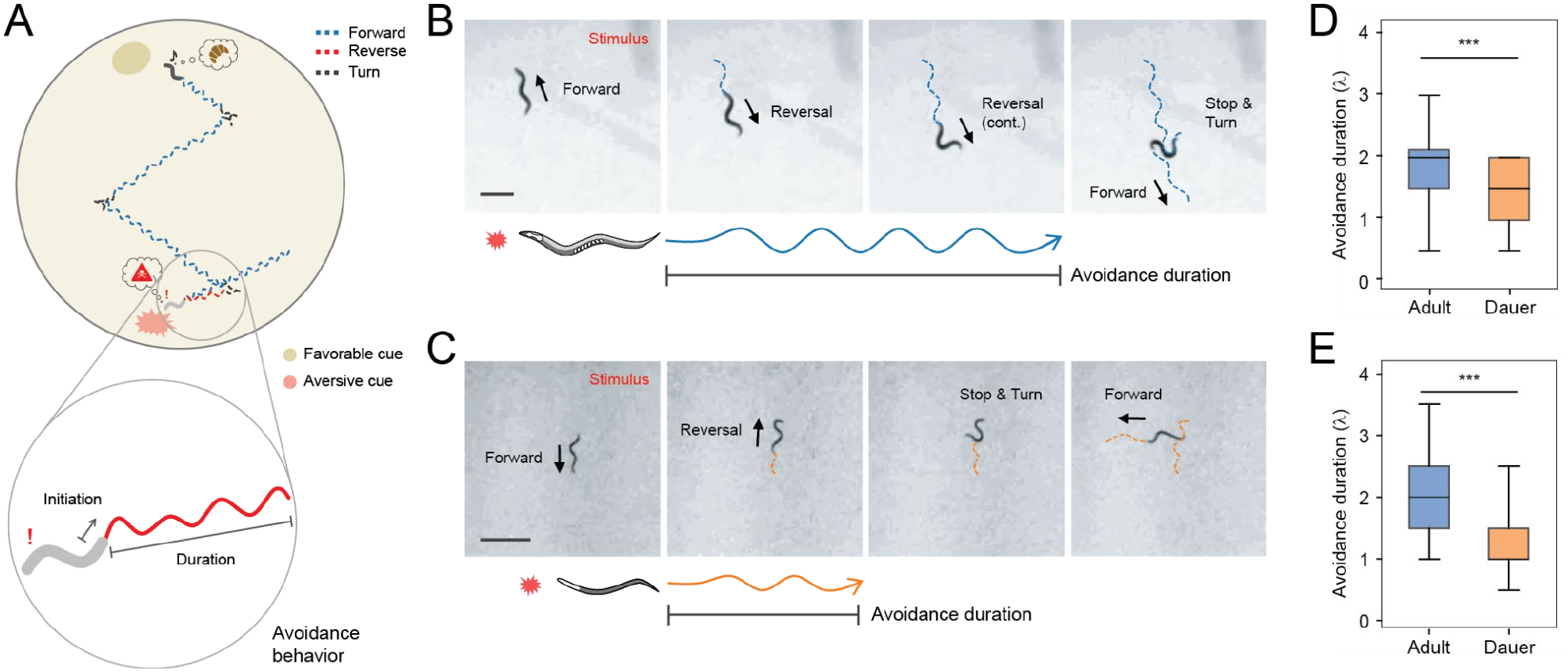
Dauers exhibit shortened avoidance duration compared with adults. **(A)** Schematic of the locomotor sequence of *C*. *elegans*. Forward, reversal, and turning movements are indicated by blue, red, and black dashed lines, respectively. Animals initiate reversals in response to aversive stimuli (red) and navigate toward favorable cues (beige). Inset, zoom-in of nociceptive escape behavior, consisting of avoidance initiation and avoidance duration. **(B,C)** Representative nociceptive escape behavior in adult (**B**) and dauer (**B**) animals with corresponding schematic representations. Red-light stimulation activates ASH sensory neurons in the first frame. Avoidance duration was defined as the interval from initiation of backward locomotion to resumption of forward movement and quantified as the number of backward sinusoidal body waves generated (blue, adult; orange, dauer). **(D,E)** Quantification of avoidance duration in adults and dauers following SDS stimulation (**D**; adults, *n* = 39; dauers, *n* = 40; \*\*\**P* = 8.3 × 10⁻⁴) and optogenetic activation of ASH neurons (**E**; adults, *n* = 36; dauers, *n* = 38; \*\*\**P* = 1.4 × 10⁻⁶). Statistics: **(D,E)** two-sided Mann–Whitney U test.

To quantify avoidance duration, we measured the number of backward sinusoidal body bends generated from reversal onset until forward locomotion resumed (*25*) (Fig. 1B,C). This metric was chosen over distance or time because dauers and adults differ in body length and locomotion speed, which would bias direct spatial or temporal comparisons. Using this normalized measure, dauers consistently produced fewer body bends during avoidance than adults, indicating that dauers terminate reversals earlier.

We exposed animals to SDS (sodium dodecyl sulfate), an aversive cue for *C*. *elegans*, and found that dauers exhibit significantly shorter avoidance duration than adults (Fig. 1D). Adults typically generated about two reversal body bends, whereas dauers showed about one (Fig. 1D). However, because stronger stimulus concentrations are required to elicit robust reversals in dauers, we asked whether the observed difference might simply reflect unequal sensory drive. To address this, we bypassed sensory input entirely by directly activating ASH neurons optogenetically and again observed a comparable reduction in avoidance duration in dauers (Fig. 1E). Furthermore, avoidance duration remained stable across a range of stimulus intensities within each stage, indicating that avoidance duration was insensitive to stimulus strength in both adults and dauers (Fig. S1A). Together, these results demonstrate that shortened avoidance duration in dauers reflects a stage-specific circuit mechanism rather than altered sensory perception.

### Dauer-specific gap junctions rewire the nociceptive circuit

Having established that dauers show shortened avoidance duration through a stage-specific circuit mechanism, we next examined how the nociceptive circuit is structurally different between the adult and dauer stages. Nociceptive circuit has been extensively studied in the context of aversive sensory processing and motor decision-making, making it an ideal substrate for investigating how dauer-specific structural rewiring shapes avoidance behavior (*26–29*). The nociceptive circuit is composed of the sensory neuron ASH and the command interneurons AVA, AVB, AVD, and PVC (*23*). Within this circuit, AVA and AVD promote backward locomotion, with AVA acting as the primary backward command interneuron, while AVB and PVC promote forward locomotion, with AVB playing the dominant role (*30*, *31*) (Fig. 2A).

**Fig. 2.**
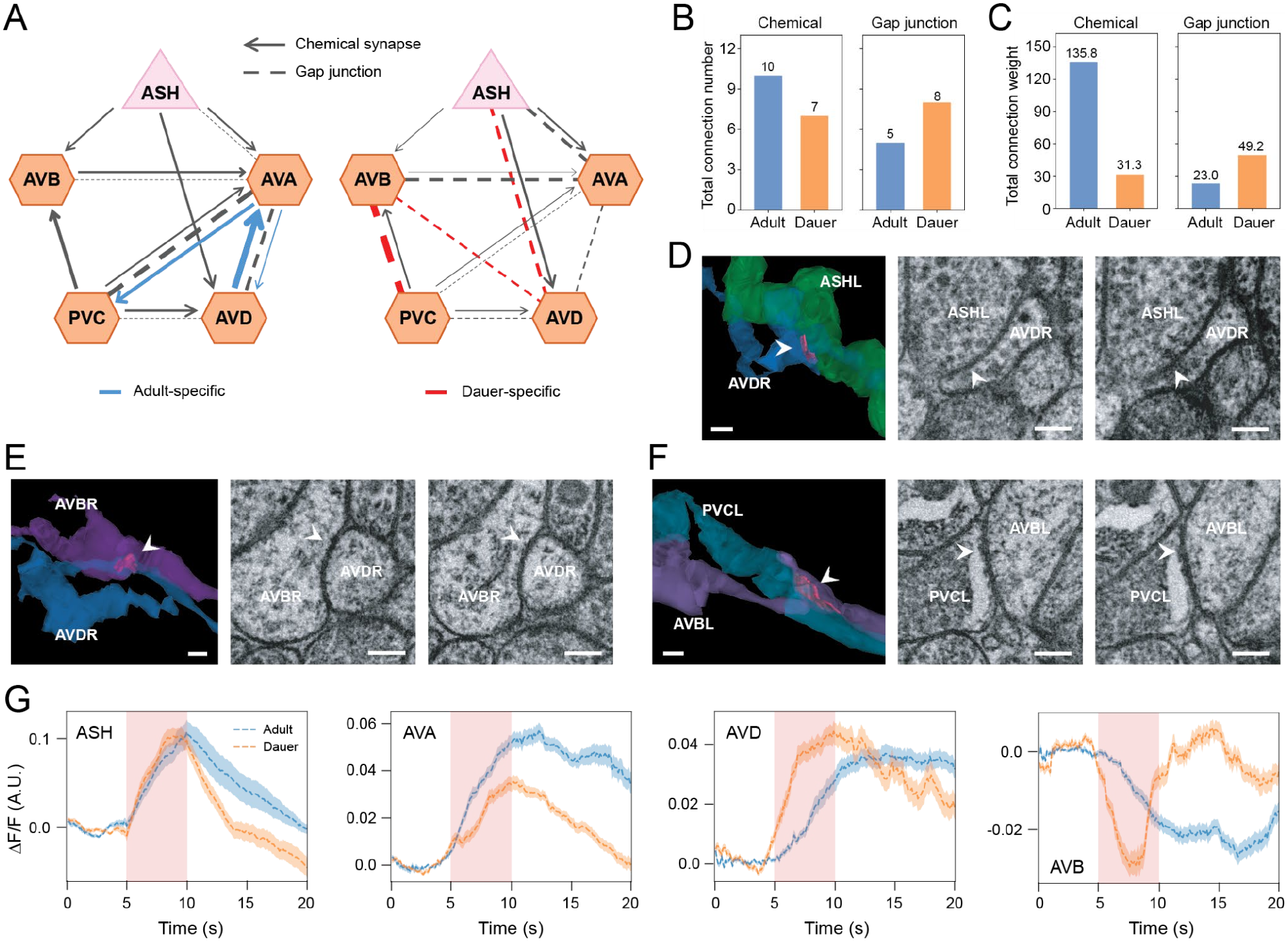
Dauer-specific electrical connections rewire the nociceptive circuit. **(A)** Nociceptive circuit diagrams in adult (left) and dauer (right) stages (adult, *2*; dauer, *4* and this study). Unless otherwise specified, “adult” refers to the adult hermaphrodite. Adult-specific connections are shown in blue and dauer-specific connections in red. Arrows indicate chemical synapses; dotted lines indicate gap junctions. The thickness of the arrow or dotted line represents the size of the synapse or gap junction (synapse weight follows Cook *et al*. (*2*) and Yim *et al*. (*4*)). Pink triangles denote sensory neurons and orange hexagons denote interneurons. **(B,C)** Quantification of total connection number (**B**) and total connection weight (**B**) of chemical synapses and gap junctions within the nociceptive circuit in adults and dauers. Connection weights were rounded to one decimal place. **(D–F)** Representative electron micrographs of dauer-specific gap junctions between ASH and AVD (**D**), AVB and AVD (**E**), and AVB and PVC (**F**), together with three-dimensional reconstructions of the corresponding contact regions. White arrowheads indicate gap junctions in the electron micrographs. Red patches in the three-dimensional reconstructions indicate gap junction contact sites. **(G)** Mean calcium activity (ΔF/F ± s.e.m.) of input neurons ASH, AVA, AVD, and the output neuron AVB during ASH stimulation. Shaded region indicates stimulus period. *n* = 15 (ASH dauer), 13 (ASH adult), 35 (AVA dauer), 24 (AVA adult), 35 (AVB dauer), 46 (AVB adult), 61 (AVD dauer), 47 (AVD adult) animals. Scale bar: 200 nm (**D–F**; 3D reconstruction), 100 nm (**D–F**; EM images).

Comparing adult and dauer nociceptive circuits extracted from published adult and dauer connectomes (*3*, *4*), we constructed circuit diagrams highlighting the stage-specific connectivity (Fig. 2A; blue: adult-specific, red: dauer-specific). To capture electrical connectivity in the dauer nervous system, we systematically annotated gap junctions, identified as parallel electron-dense membranes separated by a 2–3 nm cleft (*14*, *32–34*). These annotations were then overlaid on the previously reconstructed chemical synapse map, generating an expanded wiring diagram. As a result, we were able to find that the two stages share a common circuit backbone but differ substantially in additional gap junction connectivity (Fig. 2A). To quantify this reorganization, we compared the total number and weight of chemical synapses and gap junctions within the nociceptive circuits. Chemical connectivity was dominant in the adult stage as all the adult-specific connections were chemical synapses (Fig. 2A–C). In contrast, gap junction connectivity was markedly expanded in dauer, with both total number and connection weight significantly greater than in adults, as all the dauer-specific connections were gap junctions (Fig. 2A–C).

To confirm that these dauer-specific gap junctions represent bona fide synaptic structures, we examined the relevant contact regions in electron microscopy (EM) images. Dauer-specific electrical connections included gap junctions between ASH and AVD (ASH=AVD; hereafter “=” denotes gap junction) (Fig. 2D), AVB=AVD (Fig. 2E), and AVB=PVC (Fig. 2F). We were able to manually verify clear ultrastructural features of gap junctions—elongated, tight, and uniform apposition between neurons—in EM images for these dauer-specific gap junctions (Fig. 2D–F). These observations indicate that the dauer nociceptive circuit undergoes genuine structural remodeling through newly formed electrical synapses rather than merely reflecting differences in synapse strength. We next asked whether this rewired nociceptive circuitry reconfigures functional activities of neurons.

### Dauer-specific gap junctions reshape neuronal activity

To determine how dauer-specific circuit rewiring reconfigures neuronal activity, we performed calcium imaging of all neurons within the nociceptive circuit in both adult and dauer animals during optogenetic ASH stimulation. Both adults and dauers showed robust calcium activation in ASH and AVA during stimulation, yet dauer neurons exhibited significantly faster signal decay when the stimulus was removed (Fig. 2G). Dauer AVD neurons showed faster and more transient calcium responses when the stimulus was given (Fig. 2G). AVB, which has previously been reported to be inhibited by ASH (*23*), displayed both a faster decrease and a faster return toward baseline in dauers (Fig. 2G). PVC did not show a robust response to ASH stimulation, consistent with previous whole-brain imaging results (*35*) (data not shown).

Overall, dauer neurons showed more transient responses across all neurons in the nociceptive circuit, suggesting that the newly formed dauer-specific gap junctions accelerate signal propagation and termination within the circuit. It is interesting to note that AVB, the primary forward command interneuron, exhibited more rapid decrease and recovery from inhibition in dauers, which may directly facilitate earlier transition to forward movement following reversal behavior. We therefore asked whether dauer-specific gap junctions could produce transient calcium dynamics, leading to dauer-like avoidance behavior.

### Modeling identifies a key dauer-specific gap junction

To assess whether the calcium dynamics of dauer AVB can be explained by its connectivity-constrained presynaptic inputs, we modeled AVB activity as a weighted sum of those inputs. Based on the distinct signaling properties of electrical and chemical transmission, we modeled gap junction inputs as a fast component and chemical inputs as a slow component, resulting in a linear dynamical system with two timescales (*36*) (Fig. 3A; See Methods). Beyond the connectome-defined synapses, we included the extrasynaptic input from AVA to AVB modeled on the slow timescale (*36*) (Fig. 3B). Since PVC did not show meaningful responses during ASH stimulation, we constrained the model to three presynaptic inputs: ASH, AVA, and AVD (Fig. 3B).

**Fig. 3.**
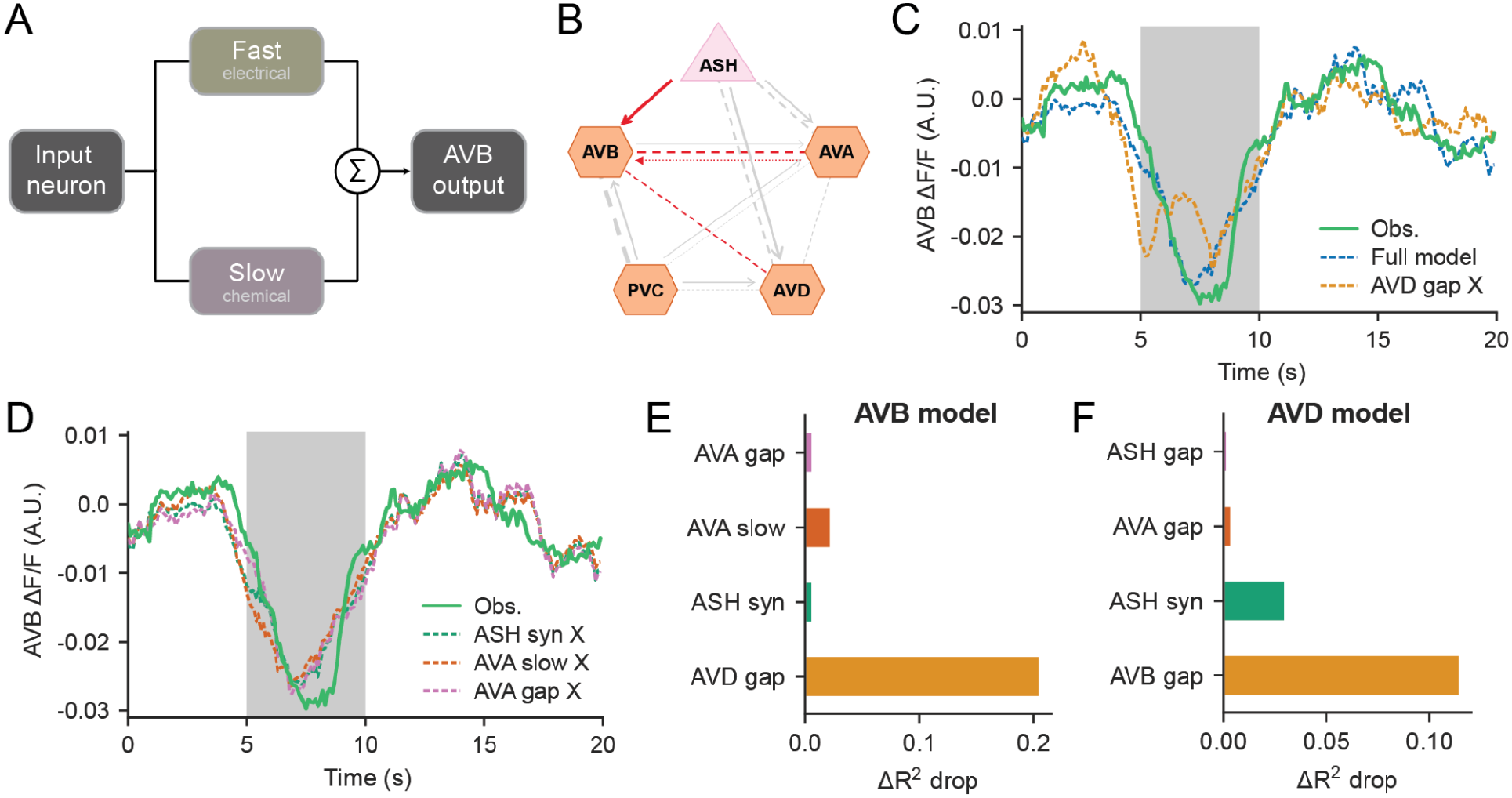
Two-timescale modeling identifies the dauer-specific AVB=AVD gap junction as a key determinant of interneuron dynamics. **(A)** Schematic of the two-timescale model. The activity of an output neuron (AVB shown) is modeled as the sum of fast (electrical) and slow (chemical or presumed extrasynaptic) components of its presynaptic inputs. **(B)** Dauer nociceptive circuit showing the connections used as inputs to the AVB model (red): chemical synapse (solid arrow), gap junctions (dashed lines), and presumed extrasynaptic connection (dotted arrow). Other connections are shown in gray. Line weight reflects connection strength. **(C)** Mean observed AVB activity in dauers (green), with fits of the full model (blue dashed; R² = 0.875) and of the model refit without the dauer-specific AVB=AVD gap junction (orange dashed; R² = 0.669). Shaded region indicates the stimulus period. **(D)** Models refit after removing connections shared between dauers and adults: the ASH chemical synapse (green), the AVA slow connection (orange), or the AVA gap junction (pink) still capture AVB activity. **(E,F)** Drop in R² (ΔR²) when each connection is removed and the remaining parameters are refit, for the AVB model **(E)** and the AVD model **(F)**. In both models, removing the AVB=AVD gap junction produced the largest drop. Models were fit to mean dauer calcium traces (*n* = 15 ASH, 35 AVA, 35 AVB, and 61 AVD animals; see Fig. 2G).

Fitting the model to the observed mean AVB calcium activity using all connected inputs yielded a strong fit (R^2^=0.875), accurately capturing both the timing of the inhibitory decrease and the subsequent recovery of AVB activity (Fig. 3C). To evaluate the specific contribution of the dauer-specific AVB=AVD, we systematically removed this connection and refitted the model. Ablation of the AVB=AVD input substantially degraded model performance (R^2^=0.669), with the model failing to precisely predict the timing of both the decay and recovery phases of AVB activity (Fig. 3C,E). Since a model incorporating all connectivity parameters could perform better simply by virtue of having more free parameters, we performed additional controls in which shared connections between dauer and adult were selectively removed. These models still captured observed AVB dynamics reasonably well (Fig. 3D,E), confirming that the model fit is not solely attributable to the number of parameters. This result indicates that the AVB=AVD is a necessary component for explaining the transient AVB dynamics observed in dauers, and cannot be compensated for by the remaining presynaptic inputs alone.

We next applied the same approach to AVD, whose dauer connectivity includes gap junctions with ASH, AVA, and AVB and a chemical synapse from ASH. Removing the AVB=AVD gap junction again produced the largest drop in model performance (ΔR² = 0.115), while removing the other connections had smaller effects (Fig. 3F). Removing the dauer-specific ASH=AVD gap junction had little effect, possibly because ASH input can also reach AVD through the ASH–AVD chemical synapse, allowing the refit model to compensate. The dauer-specific AVB=AVD gap junction therefore dominated the fits of both neurons, consistent with strong bidirectional electrical coupling between AVB and AVD in dauers.

### Synthetic gap junctions recapitulate dauer-like dynamics and behavior

To test whether the modeling results correctly predicted the importance of the dauer-specific AVB=AVD in shortening avoidance duration, we examined how introducing this connection affects neuronal activity and behavior in animals. We generated transgenic adult strains expressing a synthetic connexin-36-mediated (Cx36-mediated; (*37*)) gap junction between AVB and AVD, together with neuron-specific GCaMP6s expression in AVB and AVD, allowing direct measurement of activity changes induced by the dauer-specific AVB=AVD connection (Fig. 4A–C). Activity dynamics in both AVB and AVD shifted toward a dauer-like pattern in Cx36-expressing adults (Fig. 4D,G, Fig. S2). The AVD rise time constant following stimulus onset was significantly reduced in dauers and Cx36-expressing adults relative to wild-type adults (Fig. 4H). In AVB, the decay time constant was significantly reduced in Cx36-expressing adults to dauer-like values, while the rise time constant was not significantly changed (Fig. 4E,F). Because AVB reached similar trough levels across conditions and began recovering immediately after the trough (Fig. 4D), the overall time for AVB to recover from inhibition is determined by the sum of its decay and rise time constants. Faster decay alone was therefore sufficient to shorten AVB recovery in Cx36-expressing adults, bringing it closer to that of dauers. As AVB is the primary forward command interneuron, earlier AVB recovery provides a direct neuronal correlate for earlier resumption of forward locomotion.

**Fig. 4.**
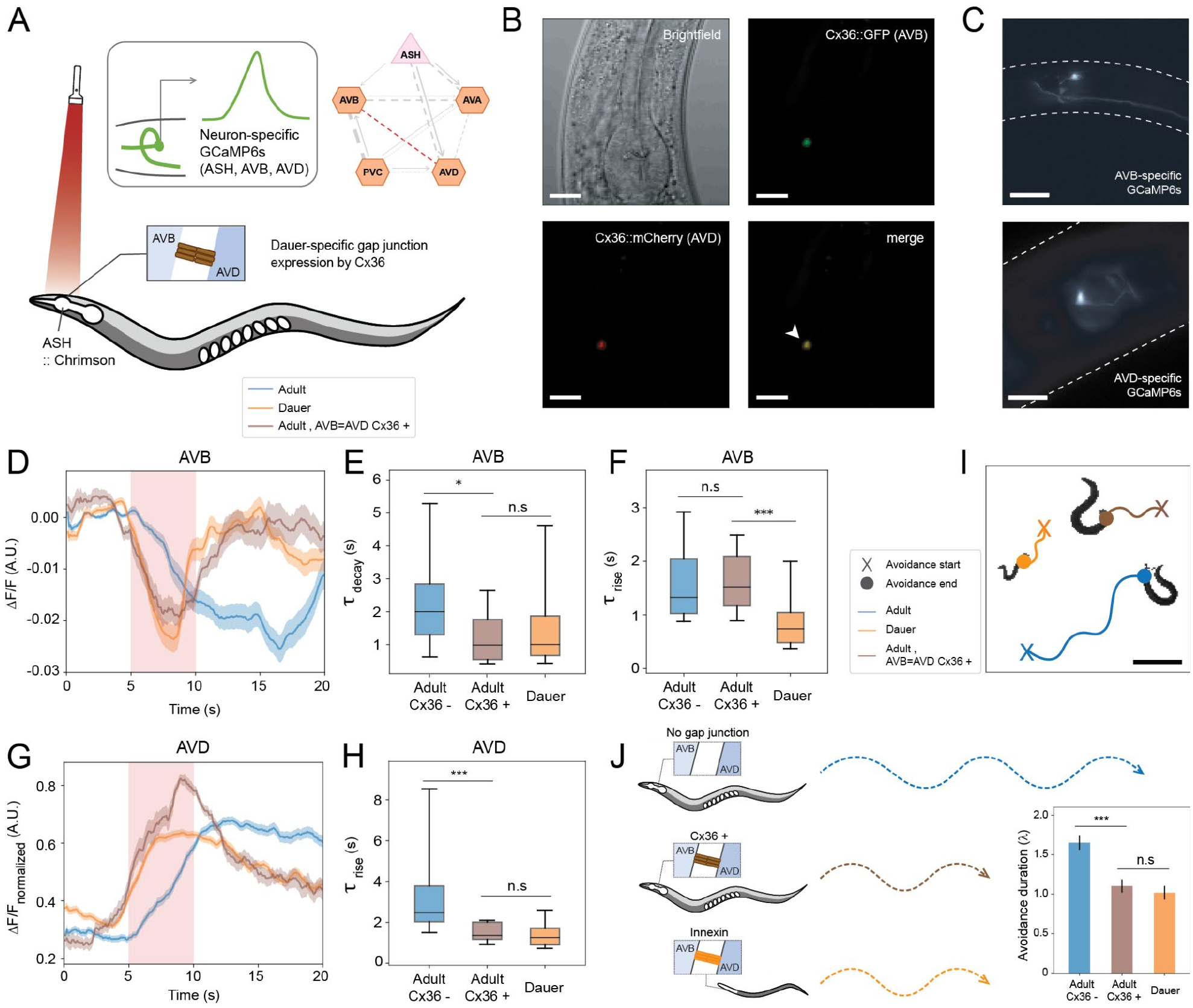
Synthetic AVB=AVD electrical coupling reproduces dauer-like circuit dynamics and behavior. **(A)** Schematic of the experimental strategy. Adult transgenic animals expressed a connexin-36 (Cx36)-mediated synthetic gap junction between AVB and AVD together with neuron-specific GCaMP6s reporters and red-light-activated channelrhodopsin (Chrimson) in ASH sensory neurons. Right, position of the dauer-specific AVB=AVD gap junction within the nociceptive circuit (red). **(B)** Confocal validation of Cx36 expression in AVB (GFP, green) and AVD (mCherry, red). Brightfield and merged fluorescence images are shown. White arrowhead indicates sites of Cx36-mediated gap junction formation (yellow in merge). **(C)** Representative images of neuron-specific GCaMP6s expression in AVB (left) and AVD (right). White dashed line outlines the worm body. **(D,G)** Calcium activity traces of AVB (**D**) and AVD (**G**) in adults (blue), dauers (orange), and adults expressing the synthetic AVB=AVD gap junction (brown). Imaging was performed at 100 ms per frame. Red shading indicates stimulation (frames 50–100; 5–10 s). Traces were smoothed using a centered moving average with a five-frame window. Sample sizes: **D**, *n* = 34 (adult Cx36−), 35 (dauer), 12 (adult Cx36+); **G**, *n* = 44 (adult Cx36−), 61 (dauer), 13 (adult Cx36+). **(E,F,H)** Comparison of response time constants in AVB (**E,F**) and AVD (**H**). **E**, decay time constant of AVB responses following stimulus onset (\**P* = 0.0124). **F**, rise time constant of AVB calcium responses measured from the stimulus-induced trough (\*\*\**P* = 0.00051). **H**, rise time constant of AVD responses following stimulus onset (\*\*\**P* = 1.6 × 10⁻⁵). **(I)** Representative locomotor trajectories during escape behavior in adults (blue), dauers (orange), and adults expressing the synthetic AVB=AVD gap junction (brown). Traces were aligned to stimulus onset and plotted at identical scale (×40). The trajectory from initiation of avoidance following stimulus delivery (marked by ‘X’) to resumption of forward locomotion (marked by ‘O’) is highlighted. Individual trajectories were recorded separately and overlaid onto a common reference frame. **(J)** Quantification of avoidance duration. Adults expressing Cx36-mediated AVB=AVD coupling exhibit shortened reversal duration comparable to dauers (*n* = 36 adult Cx36−, 34 adult Cx36+, 27 dauer; \*\*\**P* = 1.15 × 10⁻⁴). Statistics: two-sided Wilcoxon rank-sum test (**E,F,H**); two-sided Mann–Whitney U test (**J**). Scale bars: 20 µm (**B,C**), 0.5 mm (**I**). Error bars: mean ± s.e.m. (**J**).

We next asked whether these neuronal changes were sufficient to alter behavior. Adults expressing the AVB=AVD Cx36 exhibited significantly shortened avoidance duration, comparable to that observed in dauers (Fig. 4I,J). In contrast, introducing a synthetic ASH=AVA gap junction, which differs in strength between stages but is not dauer-specific, did not alter avoidance duration (Fig. S2N), indicating that the behavioral effects are specific to dauer-enriched electrical connectivity. These results support a model in which dauer-specific electrical coupling accelerates AVB recovery from inhibition, hastening the transition back to forward locomotion.

To test whether this effect generalizes across multiple dauer-specific electrical connections, we generated a second transgenic adult strain expressing a synthetic Cx36-mediated gap junction between ASH and AVD, another dauer-specific electrical connection (Fig. 5A). Although ASH decay kinetics were largely unchanged across stages and transgenic adults (Fig. 5B,C, Fig. S2), AVD responses shifted toward a dauer-like pattern (Fig. 5D, Fig. S2). The rise time constants of AVD were significantly reduced in dauers and Cx36-expressing adults relative to wild-type adults (Fig. 5E). Avoidance duration was likewise significantly shortened in Cx36-expressing adults (Fig. 5F,G), further demonstrating that dauer-specific electrical coupling is sufficient to reduce reversal duration.

**Fig. 5.**
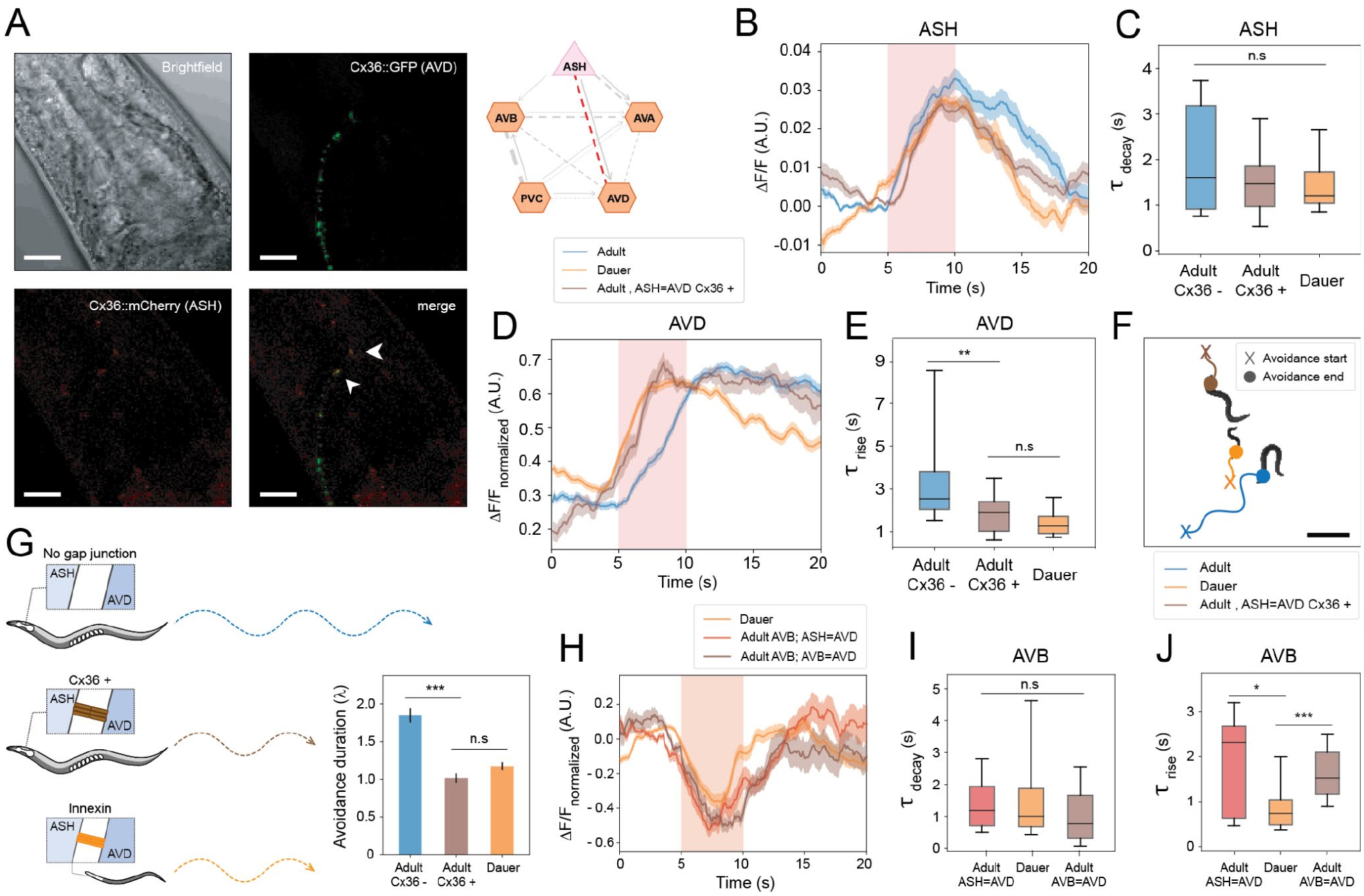
Synthetic ASH=AVD electrical coupling reproduces dauer-like escape dynamics and behavior. **(A)** Confocal validation of Cx36 expression in AVD (GFP, green) and ASH (mCherry, red). Brightfield and merged fluorescence images are shown. White arrowheads indicate sites of Cx36-mediated gap junction formation (yellow in merge). Right, position of the dauer-specific ASH=AVD gap junction within the nociceptive circuit (red). **(B,D)** Calcium activity traces of ASH (**B**) and AVD (**D**) in adults (blue), dauers (orange), and adults expressing the synthetic ASH=AVD gap junction (brown). Imaging was performed at 100 ms per frame. Red shading indicates stimulation (frames 50–100; 5–10 s). Traces were smoothed using a centered moving average with a five-frame window. Sample sizes: **B**, *n* = 11 (adult Cx36−), 12 (dauer), 18 (adult Cx36+); **D**, *n* = 44 (adult Cx36−), 61 (dauer), 13 (adult Cx36+). **(C,E)** Comparison of response time constants in ASH (**C**) and AVD (**E**). For AVD responses, rise time constants following stimulus onset differed between adult Cx36− and adult Cx36+ (\*\**P* = 0.00775). **(F)** Representative locomotor trajectories during escape behavior in adults (blue), dauers (orange), and adults expressing the synthetic ASH=AVD gap junction (brown). Traces were aligned to stimulus onset and plotted at identical scale (×40). The trajectory from initiation of avoidance following stimulus delivery (marked by ‘X’) to resumption of forward locomotion (marked by ‘O’) is highlighted. Individual trajectories were recorded separately and overlaid onto a common reference frame. **(G)** Quantification of behavioral effects of synthetic electrical coupling. Expression of a dauer-specific ASH=AVD gap junction shortened reversal duration to dauer-like levels (*n* = 30 adult Cx36−, 30 adult Cx36+, 46 dauer; \*\*\**P* = 2.7 × 10⁻⁸). **(H)** Calcium activity traces of AVB in dauers (orange), adults expressing the synthetic ASH=AVD gap junction (red), and adults expressing the synthetic AVB=AVD gap junction (brown). Imaging was performed at 100 ms per frame. Red shading indicates the stimulation period (frames 50–100; 5–10 s). Dauer and AVB=AVD Cx36+ adult traces are the same as those shown in Fig. 4D. Traces were smoothed using a centered moving average with a five-frame window. Sample sizes: *n* = 35 (dauer), 12 (adult AVB=AVD Cx36+), and 11 (adult ASH=AVD Cx36+). **(I, J)** Comparison of AVB response decay (**I**) and rise (**J**) time constants (*\*P* = 0.048; *\*\*\*P* = 0.000512). Statistics: two-sided Wilcoxon rank-sum test (**C, E, I, J**); two-sided Mann–Whitney U test (**G**). Scale bars: 20 µm (**A**) and 0.5 mm (**F**). Error bars represent mean ± s.e.m. (**G**).

Notably, dauer-like avoidance duration was reproduced without any direct electrical coupling onto the forward command interneuron AVB. We therefore asked whether the ASH=AVD connection could alter AVB activity indirectly, potentially accounting for the shortened avoidance duration. Calcium imaging revealed that AVB activity in adults expressing the synthetic ASH=AVD gap junction shifted toward a dauer-like pattern (Fig. 5H). In particular, these animals exhibited AVB signal decay comparable to that of dauers, but the AVB rise time constant remained slower than in dauers (Fig. 5I,J). Thus, a dauer-specific electrical connection can reshape AVB activity indirectly, propagating its effects through the broader circuit to alter neuronal state transitions and reduce behavioral persistence. AVB=AVD coupling likewise accelerated decay without significantly changing rise (Fig. 4E,F). Each connection therefore shortened the summed decay and rise time constants only partially, so neither connection alone fully reproduced dauer AVB kinetics (Figs. 4D and 5H). Nevertheless, in both strains AVB activity returned to baseline after stimulus onset on a time course similar to that of dauers (Figs. 4D and 5H), which likely accounts for the shortened avoidance duration to dauer-like levels in both strains (Figs. 4J and 5G). In dauers, however, the two connections may act in series. ASH=AVD shapes AVD activity, and the strong AVB=AVD coupling identified by modeling (Fig. 3E,F) relays it to AVB, so that full reconstitution of dauer AVB dynamics may require the combined action of both gap junctions.

### Avoidance initiation is preserved despite extensive circuit rewiring

While avoidance duration was strongly altered in dauers, we next asked whether avoidance initiation was similarly affected. We therefore quantified avoidance initiation using avoidance index, the fraction of trials in which an animal reverses in response to noxious stimulus, across a range of stimulus intensities (*23*) (Fig. 6A). We first simulated avoidance index curves using a previously established computational framework (*23*, *38*), adapted to the dauer circuit by scaling the surface area parameter to account for thinner dauer neurites (see Methods). Of the 667 parameter sets previously identified as valid for both hermaphrodite and male circuits, 193 also produced valid behavior in the dauer circuit. Across these sets, simulated dauer avoidance curves reached the same maximum as adult curves, with a modest rightward shift at intermediate sensory input. This shift was small compared with the difference between adult hermaphrodites and males (Fig. 6B).

**Fig. 6.**
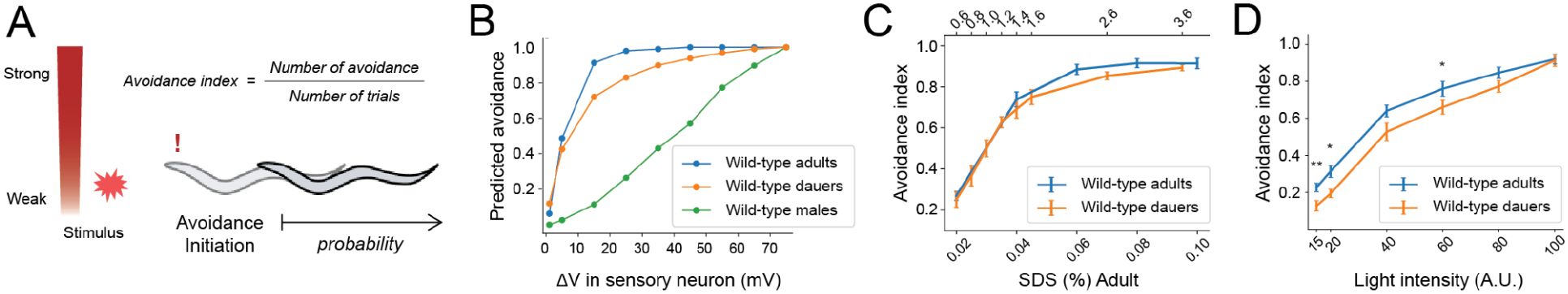
Avoidance initiation is largely preserved despite developmental circuit rewiring. **(A)** Schematic of avoidance index calculation. The avoidance index is the fraction of trials in which an animal initiates a reversal, measured across a range of stimulus intensities. **(B)** Simulated avoidance index curves for adult hermaphrodite (blue), dauer (orange), and adult male (green) nociceptive circuits as a function of the voltage change in the sensory neuron, computed across the 193 parameter sets valid for all three circuits. Predicted avoidance is the fraction of parameter sets in which A-type (backward) motor neuron activity exceeded B-type (forward) activity during stimulation. **(C)** Avoidance index measured in adults and dauers in response to SDS. Adults were tested at 0.02–0.10% SDS (bottom axis) and dauers at 0.6–3.6% SDS (top axis), as dauers require higher concentrations because of their sealed buccal cavity. The dauer axis was scaled linearly so that the lowest and highest dauer concentrations shown (0.6% and 3.6%) align with the lowest and highest adult concentrations (0.02% and 0.10%). Sample sizes (adults): *n* = 16, 13, 13, 13, and 14 for 0.02–0.10%. Sample sizes (dauers): *n* = 12, 12, 25, 22, 12, 12, 15, and 15 for 0.6–3.6%. **(D)** Avoidance index measured in adults and dauers following optogenetic activation of ASH expressing ChR2. ATR-fed animals were illuminated with blue light at the indicated intensities (*n* = 15 per group). \*\**P* = 0.0074 (15 A.U.), \**P* = 0.010 (20 A.U.), and \**P* = 0.040 (60 A.U.); *P* > 0.05 at all other intensities. Statistics: two-sided Mann–Whitney U test **(D)**. Error bars: mean ± s.e.m. **(C,D)**.

Behavioral measurements confirmed these predictions. Because dauers possess a sealed buccal cavity (*39*, *40*), substantially higher SDS concentrations were required to elicit avoidance. Over the tested ranges, dauers showed graded avoidance spanning the same range of avoidance indices as adults (Fig. 6C). To bypass the differences in chemical access, we activated ASH optogenetically. Dauers again showed graded avoidance that reached adult levels at high light intensities but was modestly lower at low intensities (Fig. 6D), consistent with the rightward shift predicted by the model. Avoidance initiation is therefore largely preserved across developmental stages despite extensive circuit rewiring (Fig. 2A–C).

## DISCUSSION

Here, we show that rewiring of the dauer nociceptive circuit selectively modifies the temporal dynamics of avoidance duration while leaving avoidance initiation largely intact (Fig. 6). Dauers terminate avoidance more rapidly and resume forward locomotion sooner than adults (Fig. 1B–E), and this behavioral difference is accompanied by distinct interneuron activity patterns that favor rapid reinstatement of forward movement (Fig. 2G). Introducing dauer-specific gap junctions into adults was sufficient to reproduce both dauer-like neuronal dynamics and shortened avoidance duration, supporting a model in which stage-specific electrical coupling contributes to the timing of circuit responses (Figs. 4 and 5).

Comparative analysis of dauer and adult connectomes further revealed an asymmetry in how the stages differ. The connections present only in adults were all chemical synapses, while those present only in dauers were all electrical synapses (Fig. 2A). At the same time, chemical synaptic connectivity is broadly weakened in the dauer circuit (Fig. 2B,C). Combined with our activity and behavioral measurements, this asymmetry suggests a developmental strategy in which electrical coupling supplements or partially replaces chemical transmission.

Introducing the dauer-specific ASH=AVD gap junction into adults also shortened avoidance duration and shifted neuronal activity toward a dauer-like state, despite the absence of a direct connection to the forward command interneuron AVB (Fig. 5H–J). This effect may be mediated through neurons and connections outside the nociceptive circuit examined here (Fig. 2A). In reproductive stage animals, transitions between forward and reverse locomotion are strongly regulated by AVA, which coordinates AVB activity across multiple timescales (*36*). The expanded gap junction connectivity in dauers may therefore shape AVB activity through both direct and indirect circuit pathways. Consistent with this, neither dauer-specific connection alone fully reconstituted the rapid recovery of dauer AVB activity. Either connection reproduced the dauer decay phase, but neither accelerated the rise phase, so each shortened AVB recovery only partially. Since both connections nonetheless shortened avoidance duration to dauer-like levels, partial acceleration of AVB recovery appears sufficient for the behavioral effect, while the full dauer AVB response may require the combined action of multiple dauer-specific gap junctions. This could be tested directly in adults carrying both connections.

Despite extensive circuit remodeling, avoidance initiation was largely preserved. Simulations predicted that dauer avoidance curves would reach adult levels with only a modest rightward shift, and behavioral measurements confirmed that dauers retained graded avoidance reaching adult levels, with modestly reduced responses only at low stimulus intensities (Fig. 6B–D). Notably, only about 30% of parameter sets that reproduce adult behavior also produced valid behavior in the dauer circuit, suggesting that the rewired dauer circuit is compatible with only part of the adult physiological regime. Since similar circuit output can arise from many combinations of intrinsic and synaptic properties (*41*), dauer neurons may undergo compensatory physiological adaptations alongside structural rewiring. Electrophysiological measurements could test whether such changes accompany dauer circuit remodeling and help preserve behavioral output.

Although motor neuron connectivity was not available in the dauer connectome, simulations incorporating alternative motor connectivity produced similar outcomes, suggesting that preserved avoidance initiation is robust to variation in downstream motor wiring (Fig. S3). Maintaining this response may be particularly important for dauer survival. Because dauers possess a sealed buccal cavity (*39*, *40*) and consequently have altered access to environmental cues, reliably converting detected aversive signals into avoidance responses may be critical.

The ability of electrical synapses to shape behavioral output is not restricted to dauer nociception. In *C*. *elegans*, electrical coupling between interneurons regulates sensory filtering and influences context-dependent behavioral choices during thermotaxis (*42*). Related roles for electrical synapses have been demonstrated in other nervous systems, including motor activity underlying stable flight in *Drosophila* (*43*), escape circuitry in zebrafish (*44*), and circadian locomotor behavior in mammals (*45*). These studies point to a shared principle across species, in which electrical synapses shape behavior by reconfiguring the temporal dynamics of neural circuits. In dauers, developmental remodeling of electrical connectivity may similarly retune circuit dynamics to adjust the persistence of an escape response while preserving its initiation.

Previous connectomic analyses showed that the overall size of the dauer chemical-synapse network at the whole-brain level remains comparable to that of adults (*4*), indicating that large-scale neural development proceeds even under developmental arrest, although chemical connectivity within the nociceptive circuit is reduced (Fig. 2B,C). Transcripts encoding gap junction components peak during early developmental stages, when electrical coupling is thought to scaffold initial circuit assembly (*46–49*) (Fig. S4). Our findings raise the possibility that dauers retain a subset of developmentally transient gap junctions that are normally pruned during maturation. Connectomes of earlier larval stages could test whether dauer-specific gap junctions overlap with early developmental electrical connections, and reconstruction of post-dauer adults could determine whether these connections are removed once reproductive development resumes.

A caveat concerns gap junction annotation. Identifying gap junctions in EM images is inherently challenging due to staining variability, oblique membrane angles, and their subtle ultrastructure (*50*, *51*), so caution is warranted when interpreting the full gap junction network at the whole-brain level. For the nociceptive circuit, we validated dauer-specific gap junctions against an independent, unpublished dataset and confirmed their presence symmetrically across left–right homologs, but similar cross-dataset confirmation will be needed for other circuits. Advances in EM imaging or staining techniques that selectively highlight gap junctions will further strengthen comparative connectomic analyses and network-level simulations (*52–55*).

In summary, similar behavioral outputs can arise from distinct internal circuit dynamics, indicating that developmental plasticity can retune how a circuit operates without altering what it computes (*41*). These findings highlight electrical synapses as versatile modulators of circuit function and establish the dauer as a model for understanding how nervous systems reconcile developmental plasticity with robust behavioral output.

## MATERIALS AND METHODS

### Strains

*C. elegans* were cultured and handled at 20°C according to standard methods (*56*). *C. elegans* Bristol strain N2 and JN2389 (peIs1090[*sra-6*p::ChR2Y2]) lines were used, obtained from the Caenorhabditis Genetics Center (CGC).

### Dauer induction

For all behavioral assays and imaging experiments, seven to twelve young adult hermaphrodites (depending on the growth rate of each strain or transgenic line) were transferred to synthetic pheromone plates to induce dauer formation. Plates contained 10 g/L agar, 7 g/L agarose, 2 g/L NaCl, 3 g/L KH₂PO₄, 0.5 g/L K₂HPO₄, 8 mg/L cholesterol, and 2 mg/L synthetic pheromone (ascaroside 2), and were seeded with *Escherichia coli* OP50. Animals were maintained at 25 °C for 4 days, after which dauers were identified based on morphological criteria, including radial body constriction and reduced body size. To ensure that only dauers were selected, plates were treated with 1.0% SDS for 30 minutes. Surviving animals were washed off with M9 buffer, transferred to unseeded plates, and allowed to recover for at least 30 minutes prior to use.

### Plasmid construction and transgenesis

All constructs generated in this study were assembled using the HiFi DNA Assembly Cloning Kit (E5520; New England Biolabs). For constructs involving the Cre/loxP system, promoters and coding sequences were cloned into the pEM1 (loxP; Addgene plasmid #24034) and pEM3 (Cre; Addgene plasmid #24033) vectors as appropriate. Promoter sequences were either amplified from N2 genomic DNA or obtained as gifts from Kyuhyung Kim and Myung-Kyu Choi. GFP and mCherry were obtained from GFP vector (pPD95.77) or mCherry vector (pPD117.01). Detailed information on assembled constructs and promoter pairs is provided in Supplementary Data.

### Transgenic strain generation

Transgenic animals were generated by microinjecting plasmid constructs into the gonads of young adult hermaphrodites using standard procedures. To isolate transgenic progeny, the following co-injection markers were used at the indicated concentrations: *unc-122*p::mCherry (25 ng/µl; red fluorescent coelomocytes), *unc-122*p::GFP (20–25 ng/µl; green fluorescent coelomocytes), *myo-2*p::mCherry (5 ng/µl; red fluorescent pharynx), and *opt-2*p::mCherry (20 ng/µl; red fluorescent intestine). Plasmid DNAs were extracted and purified using a commercial plasmid purification kit prior to injection. Full details of transgenic strains are provided in Supplementary Data.

### Fluorescence microscopy

Images of transgenic worms were acquired using a ZEISS LSM700 confocal microscope (Carl Zeiss). For imaging, worms were paralyzed with 3 mM levamisole and mounted on 3% agar pads. Confocal image stacks were processed using python to merge multiple image layers of the field of interest.

### Gap junction annotation

Gap junctions were annotated in Neuroglancer by an expert annotator blinded to cell IDs. Because gap junctions are subtle features in EM datasets, a single experienced annotator performed the entire annotation to ensure consistency. Gap junctions were marked with an annotation line connecting the two neurons involved, and the annotation was extended across all consecutive z-sections in which the junction appeared. Annotation data were then used to assign synaptic partners based on the cell IDs of the connected neurons.

Annotations that did not link two distinct neurons, or that linked a neuron to itself, were discarded. In cases where multiple annotations for the same neuronal pair were present in a given z-section and their coordinates fell within 100 nm of each other, only one annotation was retained.

### Avoidance index: SDS drop assay

Avoidance assays were performed as previously described (*22*, *23*, *57*), with minor modifications using N2 strain. Adult-stage experiments were conducted on young or 1-day-old adults, and dauer-stage experiments on 5–6 day dauers induced on pheromone plates (hereafter, “dauer” and “adult” refer to hermaphrodite dauer and hermaphrodite adult unless otherwise specified). Prior to testing, animals were transferred to foodless NGM lite plates and allowed to habituate for 10 min. Each animal then underwent 10 repellent stimulations, separated by at least 2 min.

Stimuli consisted of a small drop of sodium dodecyl sulfate (SDS) applied near the tail of a forward-moving animal using hand-pulled capillaries mounted on a mouth-controlled holder. Because dauers often remain motionless, they were gently aroused by tapping the plate until head movement was observed before stimulation. Assay plates were prepared the day before by drying unseeded NGM lite plates for 2 h at 37 °C and leaving them overnight at room temperature. Responses were scored in a 4 s window as reversal (1) or no reversal (0), and the avoidance index for each animal was calculated as the fraction of reversals across 10 trials. For adults, SDS concentrations ranged from 0.02% to 0.10%, while for dauers, concentrations from 0.6%—the lowest dose eliciting consistent responses—to 3.6% were tested.

### Avoidance index: optogenetic stimulation

Optogenetic assays were performed using strain JN2389, which expresses channelrhodopsin-2 (ChR2) in ASH neurons. This strain carries a mutation in *lite-1*, ensuring that behavioral responses reflect ChR2 activity rather than endogenous blue-light receptors. Experimental groups were raised on plates (pheromone plates for dauer induction) seeded with OP50 supplemented with 100 µM all-trans-retinal (ATR). Control groups were raised without ATR. All plates were handled in the dark to prevent ATR degradation. Young or 1-day-old adults were used for adult-stage assays, and dauers induced for 5–6 days were used for dauer-stage assays.

Animals were transferred to unseeded NGM lite plates and habituated for 10 min before testing. Experiments were performed at 22 °C using a Leica M205 FA fluorescence microscope to image and deliver light stimulation. Only animals showing forward movement before the light stimulus were used for testing. During each trial, animals were exposed to a 2 s light pulse, and reversals were scored within a 3 s window (stimulus period plus one additional second). Responses were scored as reversal (1) or no reversal (0). Each animal was tested in 10 trials with ≥2 min intervals between stimuli. Data from animals lost before completing 10 trials were excluded. The avoidance index was calculated as the mean fraction of reversals per animal.

Light intensity was varied by adjusting the laser source to six calibrated levels: 20% of 75% maximum, 20% of 100% maximum, 40% of 100% maximum, 60% of 100% maximum, 80% of 100% maximum, and 100% of 100% maximum. These corresponded to relative artificial units (A.U.) of 15, 20, 40, 60, 80, and 100, respectively.

### Avoidance duration: optogenetic stimulation

Avoidance durations were defined as the number of sinusoidal body bends executed during backward locomotion before worms resumed forward movement. To measure this under optogenetic stimulation, we used strain JN2389 or derivative transgenic lines carrying connexin-36-mediated synthetic gap junctions. The basic setup for optogenetic stimulation, including growth conditions and age-matched stages for adults and dauers, was identical to the avoidance index assay.

Animals were transferred to unseeded NGM lite plates, habituated for 10 min, and then exposed to 4 s of light stimulation. The number of sinusoidal body bends was counted manually until the animal initiated forward movement. Based on the initial posture and head orientation at stimulus onset, reversal waves were quantified using the following function:

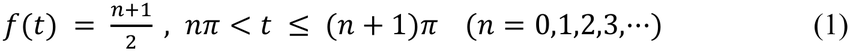

where *t* denotes how much a worm moved in sine waves. A score of 0 was assigned to animals that continued moving forward throughout stimulation or that transiently paused but resumed forward movement without executing a reversal. For measuring reversal waves in adults expressing connexin-36-mediated gap junctions, experimental and control groups were always assayed on the same day, with blind labeling performed by an independent experimenter.

Some animals expressing connexin-36-mediated synthetic gap junctions were developmentally arrested. To ensure consistent physiological comparisons, only fully developed adults carrying ectopic gap junctions—and showing no overt locomotion defects—were included in the analysis of reversal waves in adults expressing connexin-36-mediated gap junctions.

### Calcium imaging

Animals from transgenic lines in an N2 background, expressing Chrimson in ASH neurons and GCaMP6s in specific target neurons, were immobilized on 10% agarose pads prepared with a suspension of 0.10 µm microspheres (Polybead^®^, Cat# 00876-15) and sealed with a coverslip. This method rendered worms immobile throughout imaging. Calcium imaging was performed on a Nikon Eclipse Ti2-E inverted microscope equipped with a 40× water-immersion objective. Red LED light (Thorlabs, SOLIS 623C) was used for Chrimson stimulation, and blue LED light (Thorlabs, SOLIS 445C) was used for GCaMP6s excitation. Fluorescence was collected through a CFP excitation filter (Chroma Technology, AT440DC, λ = 420–442 nm) and a CFP emission filter (Chroma Technology, ET460/30x, λ = 440–480 nm). Blue light (100 mA) was continuously applied to measure baseline and inhibitory signals, while red LED stimuli (8,000 mA) were delivered manually between 5 and 10 s after recording onset for a duration of 5 s. Images were acquired at 100 ms per frame for 200 frames, yielding a total recording period of 20 s. Files in which animals moved during acquisition were discarded.

Raw nd2 files were processed using a custom Python script built on the nd2 library. For each recording, regions of interest (ROIs) were manually defined to capture neuronal somata across all frames. Fluorescence intensity was calculated as the mean pixel value within each ROI. Baseline fluorescence (F₀) was determined as the average intensity during the 50 frames (5 s) preceding stimulus onset. For each time point, *Δ*F was calculated by subtracting F₀ from the raw value, and the normalized response (*Δ*F/F₀) was obtained by dividing by F₀ to account for inter-animal variability in baseline fluorescence. All statistical analyses were performed on normalized data. For peak response comparisons, the maximum *Δ*F/F₀ value following stimulation was used; for average intensity comparisons, mean signal strength during specified time windows was analyzed.

### Calcium signal fitting

Calcium dynamics were quantified by fitting mono-exponential functions to fluorescence traces. Signal decay and rise were modeled using the following equations, respectively:

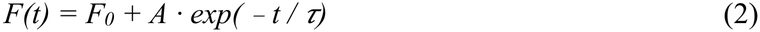

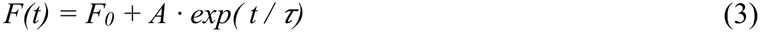

where *F(t)* denotes fluorescence at time *t*, *F_0_* is baseline fluorescence, *A* is amplitude, and *τ* is the time constant. Traces with excessive noise were smoothed by averaging over a 10-frame sliding window prior to fitting to obtain a stable estimate of *τ*. Decay constants were measured in ASH neurons following the end of a 5-s stimulus and in AVB neurons following stimulus onset. Rise constants were measured in AVA and AVD neurons following stimulus onset, and in AVB neurons following the trough of the inhibitory response. Because AVB trough amplitudes were comparable across conditions and recovery began immediately after the trough, we used the decay and rise time constants together as an index of the overall speed of AVB recovery from inhibition. Calcium signals from AVD neurons were normalized using min–max scaling prior to fitting.

### Linear dynamic modeling of interneuron activity

We modeled AVB calcium activity during ASH stimulation as a two-timescale linear function of the activity of its presynaptic input neurons. The model represents the calcium activity of the AVB neuron as a weighted sum of fast and slow components from each presynaptic input as below.

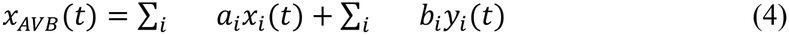

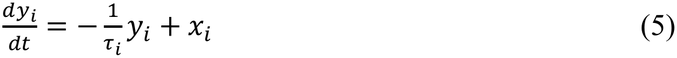

where *x_i_*(*t*) is the measured mean calcium activity of presynaptic neuron *i*, *a_i_* is the fast coupling weight representing gap junction input, *b_i_* is the slow coupling weight representing chemical synaptic or presumed extrasynaptic input, acting through a leaky-integrated copy *y_i_*(*t*) of *x_i_* with time constant *τ_i_*. Eq. 5 was discretized at the imaging frame rate (10 frames/s) using exponential decay.

Connectivity constraints were derived directly from the dauer connectome. For AVB, three presynaptic inputs were considered: ASH, AVA, and AVD. PVC was excluded as it did not show robust calcium response to ASH stimulation. Constraints were enforced as follows: ASH has no gap junction connection to AVB (*a_ASH_* = 0), and AVD has no chemical synapse onto AVB (*b_AVD_* = 0), yielding six free parameters (*a_AVA_*, *a_AVD_*, *b_ASH_*, *τ_ASH_*, *b_AVA_*, *τ_AVA_*).

Model parameters were estimated by minimizing the mean squared error between the predicted and observed mean calcium traces using differential evolution optimization. Weights were bounded to [−2, 2], and time constants to [10, 100] frames (1–10 s). Because differential evolution can converge to different local minima depending on its random seed, each model was fit with 10 different seeds, and the fit with the lowest error was kept. Model performance was evaluated using the coefficient of determination (R²).

To assess the specific contribution of individual connections, we performed systematic ablation analysis by setting the coupling weight of each connection to zero and refitting the remaining parameters. The contribution of each connection was quantified as the drop in R² (ΔR²) relative to the full connectome-constrained model. We tested ablation of the AVD gap junction, AVA gap junction, ASH chemical synapse, and AVA slow connection. AVD activity was modeled in the same way as AVB activity. Based on the dauer connectome, the AVD model included four inputs: gap junctions with ASH, AVA, and AVB, and a chemical synapse from ASH. PVC was again excluded because it showed no robust response to ASH stimulation.

### Chemical connectivity for network simulations

Original connectivity data for adult hermaphrodites and males were taken from Cook *et al*. (*2*), consistent with the datasets used in Pechuk *et al*. (*23*). For dauers, chemical synapse strengths were originally quantified differently from the adult dataset and thus required adjustment. To reconcile the two, we used the normalized adult chemical synapse connectivity matrix from Yim *et al*. (*4*), which was processed by averaging synaptic connections between left–right homologous neurons and treating them as a single cell. Scalar factors were then calculated for each connection as follows:

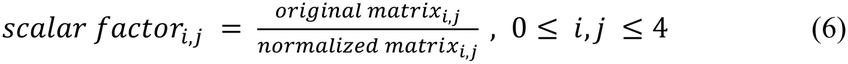

Of the 13 scalar values obtained, six were identical, each with a value of 4.755 (rounded to the fourth decimal place). This factor was therefore applied uniformly to the dauer chemical synapse connectivity matrix for all network simulations. (Here, 0 ≤ *i,j* ≤ 4, index a 5 × 5 connectivity matrix representing the neurons ASH, AVA, AVD, AVB, and PVC.) Connectivity between interneurons and A- and B-type motor neurons was adopted from the original adult hermaphrodite connectome. To test the effect of altered inter-to-motor neuron wiring in the dauer network, corresponding connections were replaced with those from the adult male connectome.

### Gap junction connectivity for network simulations

Connection weights between neuron pairs were calculated as the sum of all synaptic sizes observed between them. For gap junctions, connection strength was quantified as the number of consecutive z-sections in which the junction appeared, following the same approach used to assign connection weights in the adult connectome. All gap junction weights were incorporated into the 180 × 180 common connectivity matrix framework previously established in Yim *et al*. (*4*). For comparative analyses of the nociceptive circuit, gap junctions between left–right homologous neurons were averaged and treated as a single connection, yielding a 5 × 5 connectivity matrix consistent with that used for chemical synapses. These processed matrices were then used as input for the simulations described below. Gap junction connectivity between interneurons and A- and B-type motor neurons was likewise adopted from the adult hermaphrodite connectome. To test the effect of altered inter-to-motor neuron wiring in the dauer network, these connections were replaced with the corresponding connectivity from the adult male connectome.

### Network simulations

All network simulations were performed using the pipeline developed by Pechuk *et al*. (*23*), with minor modifications to incorporate the dauer nociceptive circuit. Because connectivity to motor neurons was not included in the dauer dataset, adult-stage motor neuron connectivity was substituted for both chemical synapses and gap junctions, yielding a 7 × 7 connectivity matrix. To assess the impact of alternative motor neuron connectivity, we additionally substituted the corresponding male motor neuron connectivity for both connection types.

Parameter sets were drawn from the 667 valid combinations identified by Pechuk *et al*. (*23*) from 15,647,317 (7^7^ × 19) tested combinations; these 667 satisfied both behavioral and physiological constraints in adult hermaphrodites and males (a kind gift from Meital Oren-Suissa). Because dauer neurons exhibit reduced neurite diameters, we adjusted the fixed surface area parameter based on average surface area values calculated from three-dimensional mesh and skeleton data derived from EM reconstructions (Adult: 15 × 10⁻⁶ cm²; Dauer: 12 × 10⁻⁶ cm²). To evaluate applicability to the dauer circuit, we applied the same behavioral and physiological constraints to the dauer network, yielding 193 valid parameter sets. These validated parameter sets were used for all subsequent simulations and analyses. For the extended membrane resistance analysis in the dauer circuit, we evaluated 4,669 parameter sets (667 × 7), corresponding to seven possible membrane resistance values ranging from 15 to 1,500 kΩ·cm², equally spaced on a logarithmic scale. This analysis yielded 1,434 valid parameter sets for the dauer network.

### Quantification and statistical analysis

All statistical analyses were performed using custom Python scripts built with the SciPy library. Two-sided Mann–Whitney U tests were used to compare distributions of reversal durations (Fig. 1, Fig. 4, Fig. 5) and two-sided Wilcoxon rank-sum tests were used to compare distributions of time constants and simulated parameters (Fig. 4, Fig. 5, Fig. 6). Statistical analyses for reversal durations were restricted to animals that exhibited a reversal response. Sample sizes and exact *P* values are reported in the corresponding figure legends.

## Acknowledgments

### Funding

This research was supported by the Samsung Science and Technology Foundation Project No. SSTF-BA1501-52 (D.T.C., J.A.B., J.L.)

AI-Bio Research Grant 0409-20230153 (D.T.C., J.A.B., J.L.)

National Research Foundation of Korea (NRF) grant 2019R1A6A1A10073437 (J.A.B.) NIH grant OD010943 (D.H.H.)

### Author contributions

Conceptualization: D.T.C., J.A.B.

Methodology: D.T.C., J.A.B.

Investigation: D.T.C., J.A.B.

Visualization: D.T.C., D.H.H., K.C.Q.N., M.C.

Supervision: J.A.B., J.L.

Writing—original draft: D.T.C., J.A.B.

Writing—review & editing: D.T.C., J.A.B., J.L.

We thank Vladyslava Pechuk, Gal Goldman, Yehuda Salzberg, and Meital Oren-Suissa for their foundational work and for providing valuable resources. We thank Hyunsoo Yim for proofreading and assisting with the assignment of gap junction annotations. Some strains were provided by the CGC, which is funded by NIH Office of Research Infrastructure Programs (P40 OD010440), and plasmid constructs were generously shared by Myung-kyu Choi and Kyuhyung Kim. We acknowledge WormBase for its indispensable data resources. We also thank Gwanho Ko and Jongmin Yoon for their help in establishing conditions for calcium imaging.

### Data, code, and materials availability

As noted in the Limitations section, certain gap junction annotations of interest require further cross-validation. For this reason, the complete dataset of dauer gap junction connectivity is not included with this publication. However, preliminary annotated electrical connectivity data are available from the corresponding author upon request. All analysis code used in this study is publicly available at https://github.com/conncslab/dauer-nociception.

## Supplementary Materials

**Fig. S1.**
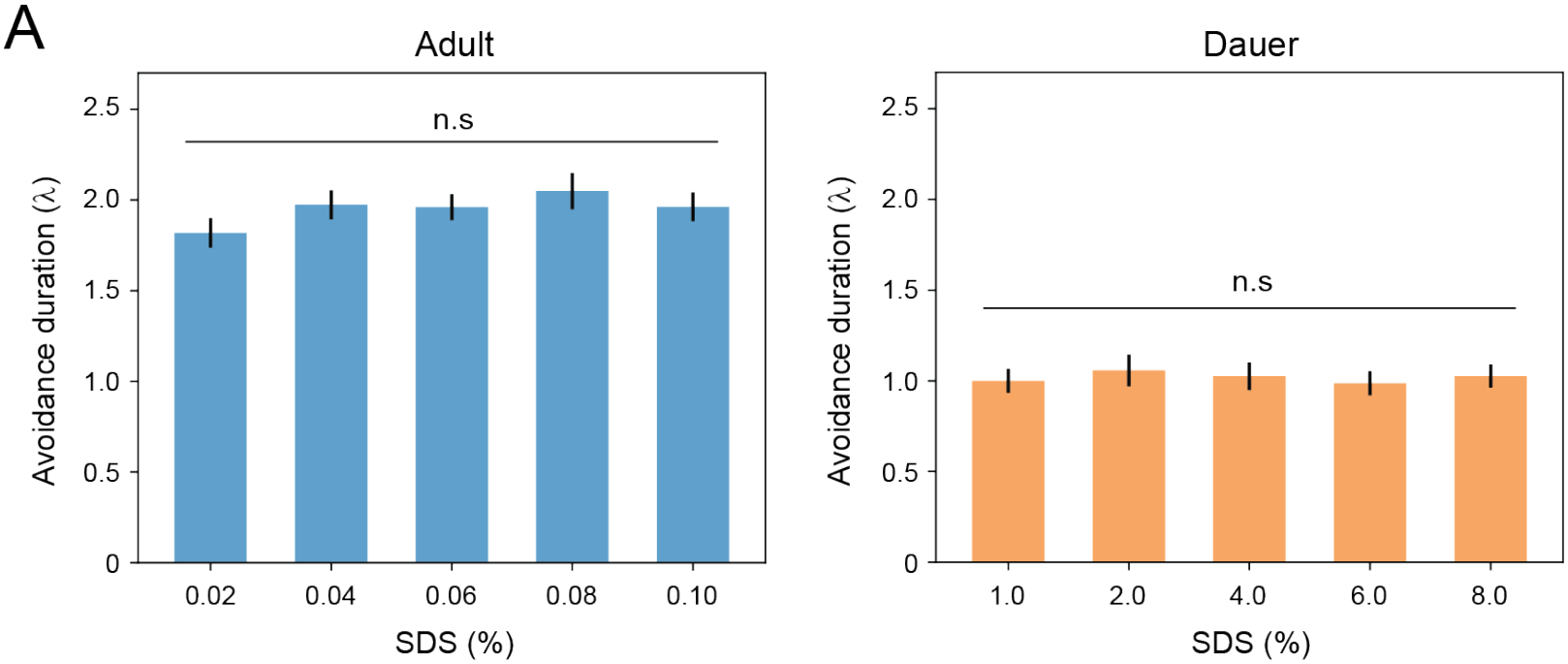
Avoidance duration is invariant across stimulus strengths within each developmental stage. **(A)** Avoidance duration across stimulus strengths for adults (left; *n* = 33, 37, 38, 41, 39 for 0.02–0.10% SDS) and dauers (right; *n* = 40, 35, 39, 38, 39 for 1.0–8.0% SDS). Statistics: **(A)** two-sided Mann–Whitney U test.

**Fig. S2.**
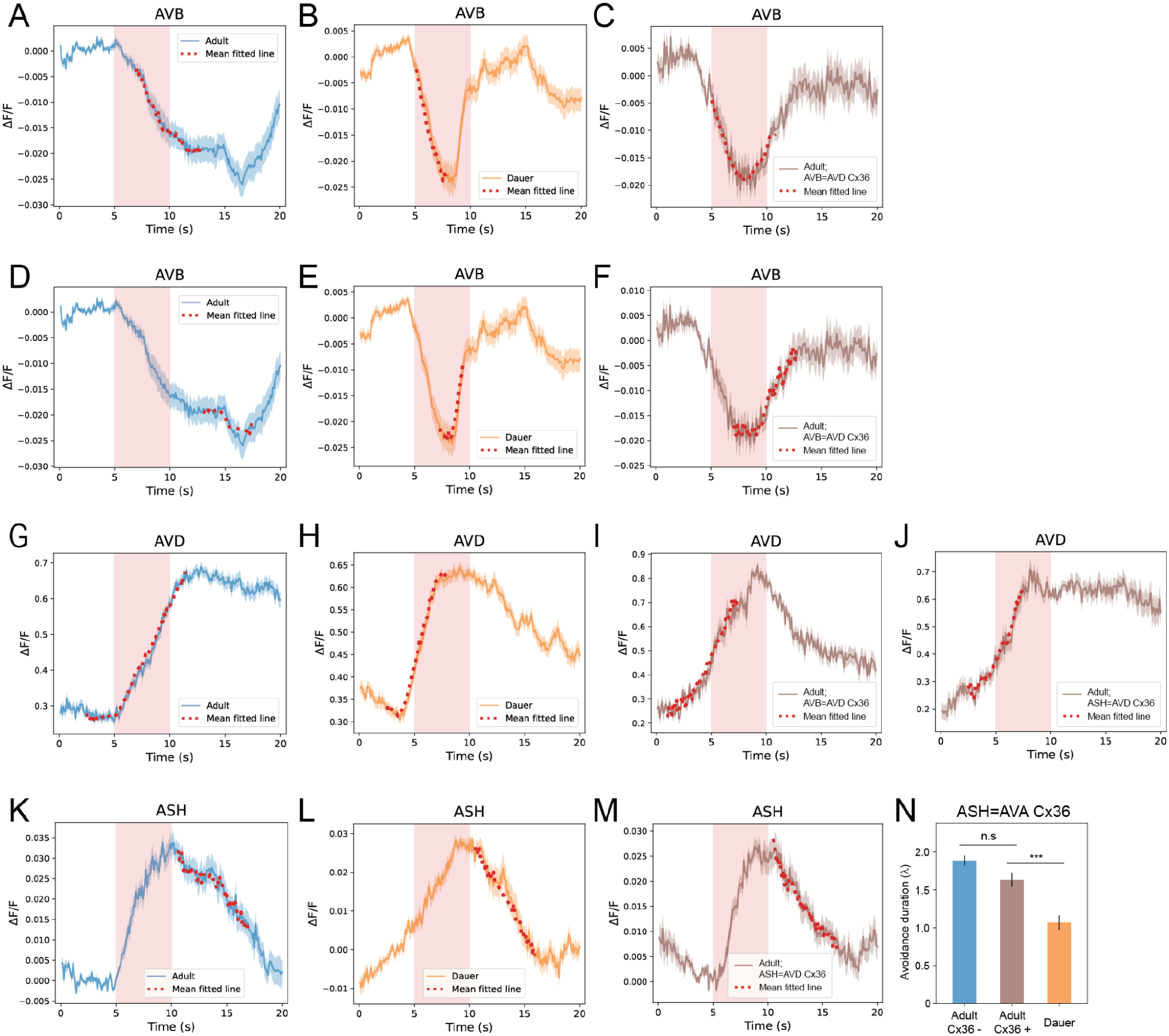
Fitted calcium response kinetics across developmental stages and synthetic gap junction conditions. Calcium activity traces and their corresponding mean fitted curves (red dotted lines) for response decay or rise. Mean fitted curves are displayed only over intervals in which at least 40% of individual traces have valid fits. **(A–C)** Calcium activity traces and mean fitted curves for AVB response decay in adults (**A**, *n* = 34), dauers (**B**, *n* = 35), and adults expressing the synthetic AVB=AVD gap junction (**C**, *n* = 12). **(D–F)** Calcium activity traces and mean fitted curves for AVB response rise in adults (**D**, *n* = 25), dauers (**E**, *n* = 34), and adults expressing the synthetic AVB=AVD gap junction (**F**, *n* = 12). Nine adult traces and one dauer trace were excluded from fitting because they did not exhibit a detectable signal rise within the 20-s recording period. **(G–J)** Calcium activity traces and mean fitted curves for AVD response rise in adults (**G**, *n* = 44), dauers (**H**, *n* = 61), adults expressing the synthetic AVB=AVD gap junction (**I**, *n* = 13), and adults expressing the synthetic ASH=AVD gap junction (**J**, *n* = 13). **(K–M)** Calcium activity traces and mean fitted curves for ASH response decay in adults (**K**, *n* = 11), dauers (**L**, *n* = 12), and adults expressing the synthetic ASH=AVD gap junction (**M**, *n* = 18). **(N)** Quantification of avoidance duration. Adults expressing shared ASH=AVA gap junction did not shorten reversal duration (*n* = 57 adult Cx36−, 57 adult Cx36+, 29 dauer; \*\*\**P* = 5.34 × 10⁻⁴). Statistics: two-sided Mann–Whitney U test **(N)**. Error bars: mean ± s.e.m. **(N)**.

**Fig. S3.**
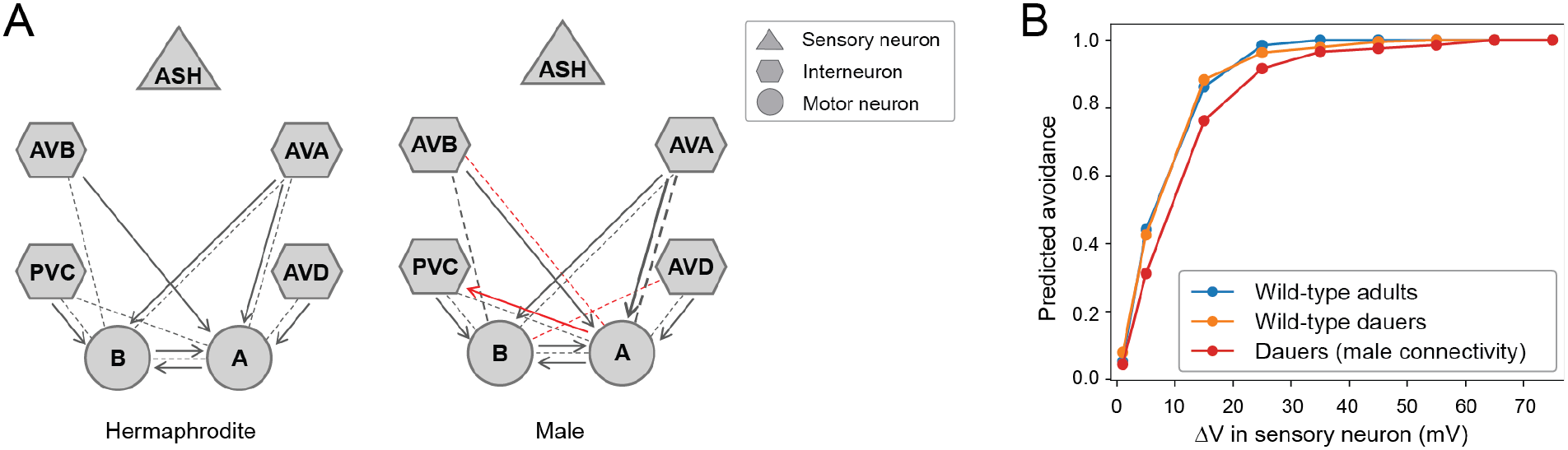
Avoidance initiation is preserved despite altered motor circuit connectivity. **(A)** Dimorphic connectivity between interneurons and motor neurons in hermaphrodite and male circuits. Dimorphic connections are highlighted in red. **(B)** Predicted avoidance index for the dauer circuit with male motor neuron connectivity.

**Fig. S4.**
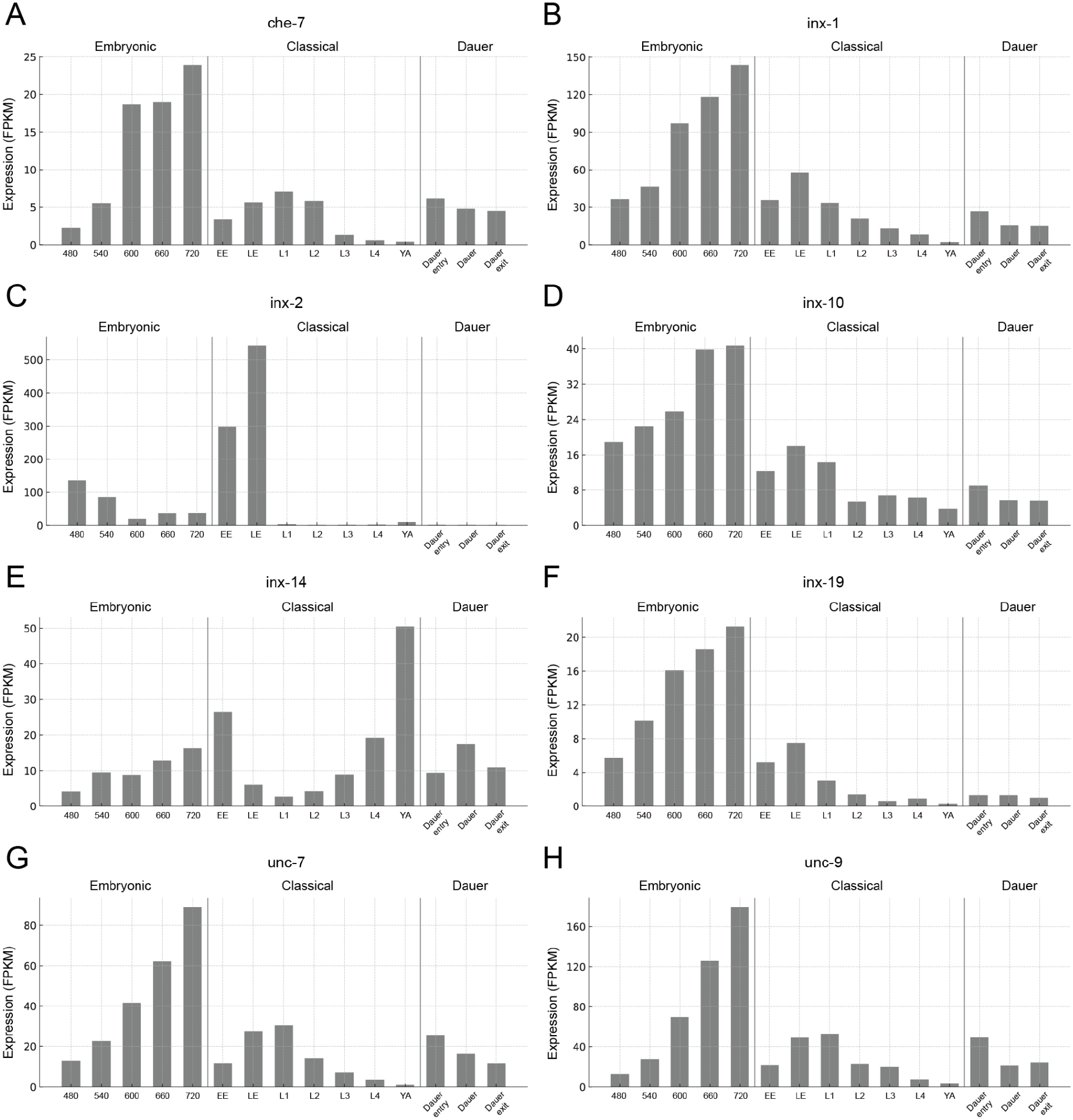
Innexin expression throughout development. **(A–H)** Developmental expression profiles of innexin genes. Bar plots show median expression levels (FPKM) of (**A**) *che-7*, (**B**) *inx-1*, (**C**) *inx-2*, (**D**) *inx-10*, (**E**) *inx-14*, (**F**) *inx-19*, (**G**) *unc-7*, and (**H**) *unc-9* across developmental stages (embryonic stage, classical stage, and dauer stage). These genes were selected as examples expressed in ASH, AVB, or AVD neurons. Expression values are from WormBase annotations.

## REFERENCES

1. J. G. White, E. Southgate, J. N. Thomson, S. Brenner, The structure of the nervous system of the nematode Caenorhabditis elegans. Philos. Trans. R. Soc. Lond. B Biol. Sci. 314, 1–340 (1986).

2. S. J. Cook, T. A. Jarrell, C. A. Brittin, Y. Wang, A. E. Bloniarz, M. A. Yakovlev, K. C. Q. Nguyen, L. T.-H. Tang, E. A. Bayer, J. S. Duerr, H. E. Bülow, O. Hobert, D. H. Hall, S. W. Emmons, Whole-animal connectomes of both Caenorhabditis elegans sexes. Nature 571, 63–71 (2019).

3. D. Witvliet, B. Mulcahy, J. K. Mitchell, Y. Meirovitch, D. R. Berger, Y. Wu, Y. Liu, W. X. Koh, R. Parvathala, D. Holmyard, R. L. Schalek, N. Shavit, A. D. Chisholm, J. W. Lichtman, A. D. T. Samuel, M. Zhen, Connectomes across development reveal principles of brain maturation. Nature 596, 257–261 (2021).

4. H. Yim, D. T. Choe, J. A. Bae, M.-K. Choi, H.-M. Kang, K. C. Q. Nguyen, S. Ahn, S.-K. Bahn, H. Yang, D. H. Hall, J. S. Kim, J. Lee, Comparative connectomics of dauer reveals developmental plasticity. Nat Commun 15, 1546 (2024).

5. R. C. Cassada, R. L. Russell, The dauerlarva, a post-embryonic developmental variant of the nematode Caenorhabditis elegans. Dev. Biol. 46, 326–342 (1975).

6. J. W. Golden, D. L. Riddle, The Caenorhabditis elegans dauer larva: developmental effects of pheromone, food, and temperature. Dev Biol 102, 368–378 (1984).

7. E. A. Hallem, A. R. Dillman, A. V. Hong, Y. Zhang, J. M. Yano, S. F. DeMarco, P. W. Sternberg, A sensory code for host seeking in parasitic nematodes. Curr Biol 21, 377– 383 (2011).

8. E. A. Hallem, P. W. Sternberg, Acute carbon dioxide avoidance in Caenorhabditis elegans. Proc. Natl. Acad. Sci. U. S. A. 105, 8038–8043 (2008).

9. A. J. Bretscher, K. E. Busch, M. de Bono, A carbon dioxide avoidance behavior is integrated with responses to ambient oxygen and food in Caenorhabditis elegans. Proc. Natl. Acad. Sci. U. S. A. 105, 8044–8049 (2008).

10. N. Fielenbach, A. Antebi, C. elegans dauer formation and the molecular basis of plasticity. Genes Dev. 22, 2149–2165 (2008).

11. H. Lee, M.-K. Choi, D. Lee, H.-S. Kim, H. Hwang, H. Kim, S. Park, Y.-K. Paik, J. Lee, Nictation, a dispersal behavior of the nematode Caenorhabditis elegans, is regulated by IL2 neurons. Nat. Neurosci. 15, 107–112 (2011).

12. J. A. Bae, M.-K. Choi, S. Ahn, G. Ko, D. T. Choe, H. Yim, K. C. Nguyen, J. S. Kim, D. H. Hall, J. Lee, Structural diversity of mitochondria in the neuromuscular system across development revealed by 3D electron microscopy. Adv. Sci. (Weinh.) 12, e2411191 (2025).

13. A. E. Pereda, Electrical synapses and their functional interactions with chemical synapses. Nat. Rev. Neurosci. 15, 250–263 (2014).

14. D. A. Goodenough, D. L. Paul, Gap junctions. Cold Spring Harb. Perspect. Biol. 1, a002576 (2009).

15. M. V. L. Bennett, R. S. Zukin, Electrical coupling and neuronal synchronization in the Mammalian brain. Neuron 41, 495–511 (2004).

16. P. Alcamí, A. E. Pereda, Beyond plasticity: the dynamic impact of electrical synapses on neural circuits. Nat. Rev. Neurosci. 20, 253–271 (2019).

17. J. Song, K. Ampatzis, E. R. Björnfors, A. El Manira, Motor neurons control locomotor circuit function retrogradely via gap junctions. Nature 529, 399–402 (2016).

18. O. Kiehn, M. C. Tresch, Gap junctions and motor behavior. Trends Neurosci. 25, 108–115 (2002).

19. D. H. Hall, Gap junctions in C. elegans: Their roles in behavior and development: Gap Junctions in*C. elegans*. Dev. Neurobiol. 77, 587–596 (2017).

20. C. I. Bargmann, J. H. Thomas, H. R. Horvitz, Chemosensory cell function in the behavior and development of Caenorhabditis elegans. Cold Spring Harb. Symp. Quant. Biol. 55, 529–538 (1990).

21. J. M. Kaplan, H. R. Horvitz, A dual mechanosensory and chemosensory neuron in Caenorhabditis elegans. Proc. Natl. Acad. Sci. U. S. A. 90, 2227–2231 (1993).

22. M. A. Hilliard, C. I. Bargmann, P. Bazzicalupo, C. elegans responds to chemical repellents by integrating sensory inputs from the head and the tail. Curr. Biol. 12, 730–734 (2002).

23. V. Pechuk, G. Goldman, Y. Salzberg, A. H. Chaubey, R. A. Bola, J. R. Hoffman, M. L. Endreson, R. M. Miller, N. J. Reger, D. S. Portman, D. M. Ferkey, E. Schneidman, M. Oren-Suissa, Reprogramming the topology of the nociceptive circuit in C. elegans reshapes sexual behavior. Curr Biol 32, 4372–4385.e7 (2022).

24. A. C. Hart, S. Sims, J. M. Kaplan, Synaptic code for sensory modalities revealed by C. elegans GLR-1 glutamate receptor. Nature 378, 82–85 (1995).

25. J. M. Gray, J. J. Hill, C. I. Bargmann, A circuit for navigation in Caenorhabditis elegans. Proc Natl Acad Sci U S A 102, 3184–3191 (2005).

26. M. Guo, T.-H. Wu, Y.-X. Song, M.-H. Ge, C.-M. Su, W.-P. Niu, L.-L. Li, Z.-J. Xu, C.-L. Ge, M. T. H. Al-Mhanawi, S.-P. Wu, Z.-X. Wu, Reciprocal inhibition between sensory ASH and ASI neurons modulates nociception and avoidance in Caenorhabditis elegans. Nat. Commun. 6, 5655 (2015).

27. M. C. Krzyzanowski, S. Woldemariam, J. F. Wood, A. H. Chaubey, C. Brueggemann, A. Bowitch, M. Bethke, N. D. L’Etoile, D. M. Ferkey, Aversive behavior in the nematode C. elegans is modulated by cGMP and a neuronal gap junction network. PLoS Genet. 12, e1006153 (2016).

28. J.-J. Wu, S.-W. Yin, H. Liu, R. Li, J.-H. Huang, P.-Z. Wang, Y. Xu, J.-L. Zhao, P.-P. Wu, Z.-X. Wu, Positive interaction between ASH and ASK sensory neurons accelerates nociception and inhibits behavioral adaptation. iScience 25, 105287 (2022).

29. C. Lin, Y. Shan, Z. Wang, H. Peng, R. Li, P. Wang, J. He, W. Shen, Z. Wu, M. Guo, Molecular and circuit mechanisms underlying avoidance of rapid cooling stimuli in C. elegans. Nat. Commun. 15, 297 (2024).

30. M. Chalfie, J. E. Sulston, J. G. White, E. Southgate, J. N. Thomson, S. Brenner, The neural circuit for touch sensitivity in Caenorhabditis elegans. J Neurosci 5, 956–964 (1985).

31. S. R. Wicks, C. J. Roehrig, C. H. Rankin, A dynamic network simulation of the nematode tap withdrawal circuit: predictions concerning synaptic function using behavioral criteria. J Neurosci 16, 4017–4031 (1996).

32. A. C. Miller, A. E. Pereda, The electrical synapse: Molecular complexities at the gap and beyond: Structural Complexity of Electrical Synapses. Dev. Neurobiol. 77, 562–574 (2017).

33. D. H. Hall, WormAtlas Hermaphrodite Handbook - Gap Junctions. WormAtlas, doi: 10.3908/wormatlas.1.25 (2018).

34. S. A. Bloomfield, B. Völgyi, The diverse functional roles and regulation of neuronal gap junctions in the retina. Nat. Rev. Neurosci. 10, 495–506 (2009).

35. F. Randi, A. K. Sharma, S. Dvali, A. M. Leifer, Neural signal propagation atlas of Caenorhabditis elegans. Nature 623, 406–414 (2023).

36. J. Meng, T. Ahamed, B. Yu, W. Hung, S. Ei Mouridi, Z. Wang, Y. Zhang, Q. Wen, T. Boulin, S. Gao, M. Zhen, A tonically active master neuron modulates mutually exclusive motor states at two timescales. Sci Adv 10, eadk0002 (2024).

37. I. Rabinowitch, M. Chatzigeorgiou, B. Zhao, M. Treinin, W. R. Schafer, Rewiring neural circuits by the insertion of ectopic electrical synapses in transgenic C. elegans. Nat. Commun. 5, 4442 (2014).

38. W. Gerstner, W. M. Kistler, R. Naud, L. Paninski, Neuronal Dynamics: From Single Neurons to Networks and Models of Cognition (Cambridge University Press, Cambridge, England, 2014; 10.1017/cbo9781107447615).

39. R. J. Androwski, K. M. Flatt, N. E. Schroeder, Phenotypic plasticity and remodeling in the stress-induced Caenorhabditis elegans dauer: Phenotypic plasticity and remodeling. Wiley Interdiscip. Rev. Dev. Biol. 6, e278 (2017).

40. C. Wolkow, D. H. Hall, WormAtlas Dauer Handbook - The Dauer Cuticle. WormAtlas, doi: 10.3908/wormatlas.3.1 (2011).

41. A. A. Prinz, D. Bucher, E. Marder, Similar network activity from disparate circuit parameters. Nat. Neurosci. 7, 1345–1352 (2004).

42. A. Almoril-Porras, A. C. Calvo, L. Niu, J. Beagan, M. Díaz García, J. D. Hawk, A. Aljobeh, E. M. Wisdom, I. Ren, Z.-W. Wang, D. A. Colón-Ramos, Configuration of electrical synapses filters sensory information to drive behavioral choices. Cell 188, 89– 103.e13 (2025).

43. S. Hürkey, N. Niemeyer, J.-H. Schleimer, S. Ryglewski, S. Schreiber, C. Duch, Gap junctions desynchronize a neural circuit to stabilize insect flight. Nature 618, 118–125 (2023).

44. A. C. Miller, A. C. Whitebirch, A. N. Shah, K. C. Marsden, M. Granato, J. O’Brien, C. B. Moens, A genetic basis for molecular asymmetry at vertebrate electrical synapses. Elife 6, e25364 (2017).

45. M. A. Long, M. J. Jutras, B. W. Connors, R. D. Burwell, Electrical synapses coordinate activity in the suprachiasmatic nucleus. Nat. Neurosci. 8, 61–66 (2005).

46. Alliance of Genome Resources Consortium, Updates to the Alliance of Genome Resources central infrastructure. Genetics 227, iyae049 (2024).

47. J.-W. Cao, L.-Y. Liu, Y.-C. Yu, Gap junctions regulate the development of neural circuits in the neocortex. Curr. Opin. Neurobiol. 81, 102735 (2023).

48. K. Kandler, L. C. Katz, Neuronal coupling and uncoupling in the developing nervous system. Curr. Opin. Neurobiol. 5, 98–105 (1995).

49. L. A. B. Elias, A. R. Kriegstein, Gap junctions: multifaceted regulators of embryonic cortical development. Trends Neurosci. 31, 243–250 (2008).

50. M. Pallotto, P. V. Watkins, B. Fubara, J. H. Singer, K. L. Briggman, Extracellular space preservation aids the connectomic analysis of neural circuits. Elife 4, e08206 (2015).

51. G. E. Sosinsky, B. J. Nicholson, Structural organization of gap junction channels. Biochim. Biophys. Acta 1711, 99–125 (2005).

52. M. Ishibashi, J. Keung, C. W. Morgans, S. A. Aicher, J. R. Carroll, J. H. Singer, L. Jia, W. Li, I. Fahrenfort, C. P. Ribelayga, S. C. Massey, Analysis of rod/cone gap junctions from the reconstruction of mouse photoreceptor terminals. Elife 11, e73039 (2022).

53. J. S. Goldberg, T. J. Vadakkan, K. K. Hirschi, M. E. Dickinson, A computational approach to detect gap junction plaques and associate them with cells in fluorescent images. J. Histochem. Cytochem. 61, 283–293 (2013).

54. S. M. Markert, S. Britz, S. Proppert, M. Lang, D. Witvliet, B. Mulcahy, M. Sauer, M. Zhen, J.-L. Bessereau, C. Stigloher, Filling the gap: adding super-resolution to array tomography for correlated ultrastructural and molecular identification of electrical synapses at the C. elegans connectome. Neurophotonics 3, 041802 (2016).

55. B. Mulcahy, D. Witvliet, D. Holmyard, J. Mitchell, A. D. Chisholm, Y. Meirovitch, A. D. T. Samuel, M. Zhen, A pipeline for volume electron microscopy of the Caenorhabditis elegans nervous system. Front. Neural Circuits 12, 94 (2018).

56. S. Brenner, The genetics of Caenorhabditis elegans. Genetics 77, 71–94 (1974).

57. M. Oren-Suissa, E. A. Bayer, O. Hobert, Sex-specific pruning of neuronal synapses in Caenorhabditis elegans. Nature 533, 206–211 (2016).

